# The Lateral/Caudal Ganglionic Eminence Makes a Limited Contribution to Cortical Oligodendrocytes

**DOI:** 10.1101/2023.12.01.569674

**Authors:** Jialin Li, Feihong Yang, Yu Tian, Ziwu Wang, Dashi Qi, Zhengang Yang, Jiangang Song, Jing Ding, Xin Wang, Zhuangzhi Zhang

## Abstract

The emergence of myelinating oligodendrocytes represents a pivotal developmental milestone in vertebrates, given their capacity to ensheath axons and facilitate the swift conduction of action potentials. It is widely accepted that cortical oligodendrocyte progenitor cells (OPCs) arise from medial ganglionic eminence (MGE), lateral/caudal ganglionic eminence (LGE/CGE) and cortical radial glial cells (RGCs). Here, we used two different fate mapping strategies to challenge the established notion that the LGE generates cortical OPCs. Furthermore, we used a Cre/loxP-dependent exclusion strategy to reveal that the LGE/CGE-derived OPCs are minimal. Additionally, we showed that specifically eliminating MGE-derived OPCs leads to a significant reduction of cortical OPCs. Together, our findings indicate that the contribution of OPCs from LGE/CGE is minimal, contrary to previous beliefs. These findings provide a new view of the developmental origins of cortical OPCs and a valuable foundation for future research on both normal development and oligodendrocyte-related disease.

## Introduction

Oligodendrocytes (OLs) are one of the most abundant cell types in the mammalian neocortex. The emergence of myelination OLs in vertebrates is a major event in the evolution of the central nervous system (CNS), as these cells play a crucial role in enhancing axonal insulation and the conduction of neural impulses. Previously, they have been described as support cells that myelinate axons and play essential roles in the function of neocortical circuits. Myelin loss or dysfunction is associated with many disorders, such as multiple sclerosis, depression and pediatric leukodystrophies^1,2^.

During CNS development, OLs originate from oligodendrocyte precursor cells (OPCs), which arise from different regions of the embryonic neural tube. OPCs proliferate and migrate away from the ventricular germinal zone (VZ) into developing gray and white matter before differentiating into mature OLs^3,4^. The developmental origin of cortical oligodendrocytes has been extensively studied in the CNS, especially in the neocortex^5–8^. A landmark study^9^ demonstrated that cortical OPCs are generated in three waves, starting with a ventral wave, gradually transitioning to more dorsal origins. The first wave of cortical OPCs is derived from the medial ganglionic eminence (MGE) or the anterior entopeduncular area (AEP) at embryonic day E12.5 (*Nkx2.1*^+^ lineage). The second wave of cortical OLs is derived from the lateral/caudal ganglionic eminences (LGE/CGE) at E15.5 (*Gsx2^+^/Nkx2.1^-^* lineage). The final wave occurs at P0, when OPCs originate from cortical glial progenitor cells (*Emx1*^+^ lineage). Our recent studies^10,11^, however, challenge the notion that cortical OLs are generated in these three waves^9,12–14^, as the cortical glial progenitor cells were also found to express *Gsx2*, which was known to be exclusively expressed in the developing subpallium^15–17^, after E16.5^11^. Therefore, *Gsx2^+^/Nkx2.1^-^* lineage OLs may not represent LGE-derived cortical OLs. To date, whether the LGE/CGE generates cortical OPCs (OLs) during cortical development remains an unanswered question in the field of developmental neuroscience.

The present study was undertaken to answer this question. We carried out distinct fate mapping experiments and intersectional lineage analyses to reveal that cortical OPCs do not derive from the LGE/CGE. First, using in utero electroporation (IUE) to target the LGE in a Cre recombinase-dependent *IS* (*Rosa^CAG-LSLFrt-tdTomato-Frt-EGFP^*) reporter mouse line^18^, we found that LGE-derived OPCs produced striatal OLs rather than cortical OLs. Second, by combining the PiggyBac transposon system with IUE, we found that no traced cells migrated into the cortex from the LGE. Third, by crossing the *Nkx2.1^Cre^* and *Emx1^Cre^* mouse lines with an *Rosa^H2B-GFP^* reporter mouse line, we found that nearly all cortical OPCs originated from *Nkx2.1^Cre^* or *Emx1*^Cre^ lineage cells. Finally, we analyzed *Olig2*-NCKO (*Nkx2.1^Cre^; Olig2^F/F^*) mice, and found that the number of cortical OPCs was significantly reduced from E14.5 to E16.5. These results indicated that the MGE was the predominant ventral source of cortical OPCs. Taken together, our findings suggest that the LGE/CGE gives rise to limited cortical OPCs (OLs) during neocortical development.

## Results

### Fate mapping of LGE-derived OPCs by combining IUE with a Cre recombinase-dependent *IS* reporter

OPCs initially emerge in the MGE and can be labeled by platelet-derived growth factor receptor (a-subunit, *Pdgfra*) around E12.5. We hypothesized that if the LGE generated cortical OPCs, the *Pdgfra^+^* cell migration stream from the LGE to the cortex could be observed during cortical development. To confirm this speculation, we conducted a detailed analysis of *Pdgfra* mRNA expression using *in situ* hybridization (ISH). Our results showed that there was no observable *Pdgfra^+^* cell migration stream from the LGE to the cortex from E13.5 to P0 (Figure 1A). Next, we used Cre recombinases in combination with an *IS* reporter mouse line^18^ to fate map LGE-derived OPCs (OLs). We delivered pCAG-Cre plasmids specifically into the LGE VZ of *IS* reporter mouse embryos at E13.5 by IUE (Figures 1B-C). In this experiment, cells generated from electroporated neural stem cells (NSCs) expressed tdTomato (tdT). Interestingly, we observed many tdT^+^ cells throughout the whole LGE (striatum) at E18.5. In contrast, there were very few tdT^+^ cells detected in the cortex (Figure 1D). OPCs express *Olig2*, *Sox10*, *Pdgfra* and NG2 (*Cspg4*) and mature neurons express NeuN ^19–25^. To determine the identity of the tdT^+^ cells, we performed double staining of tdT with OLIG2, SOX10, PDGFRA, NG2 (CSPG4) and NeuN. Within the cortex, tdT^+^ cells did not express OPC marker genes but express NeuN; however, tdT^+^ cells expressed OPC marker genes and NeuN in the striatum (Figure 1E-F). We also delivered pCAG-Cre plasmids specifically into the LGE VZ of *IS* reporter mice at P0 by IUE and analyzed these mice at P49 (Figure 1G) to investigate whether the LGE generates cortical OLs at a later developmental stage. Our results showed that LGE progenitors generate striatum OLs and astrocytes, which expressed SOX10 and SOX9, respectively (Figure 1G).

**Figure 1.**
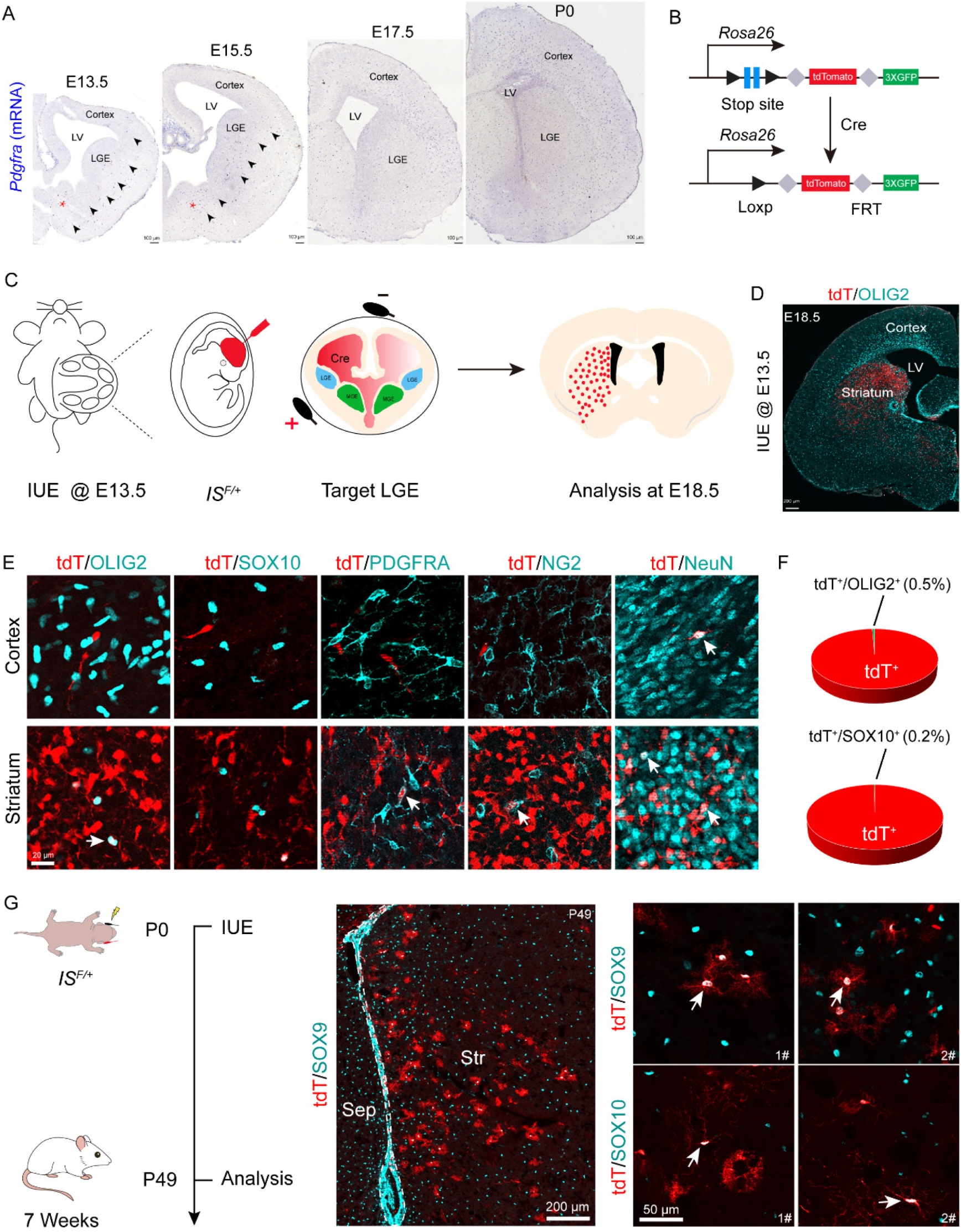
Fate mapping of LGE-derived OPCs by combining IUE with a Cre recombinase-dependent IS reporter. (A) *In situ* hybridization showing *Pdgfra* expression from E13.5 to P0. Arrowheads indicated the *Pdgfra^+^* cell migration stream from the MGE/AEP to the cortex. (B) Scheme of the IS reporter lines. (C) Experimental schedule to trace LGE-derived OPCs. (D) Representative coronal sections showing the distribution of tdT^+^ cells. (E) tdT^+^ cells expressed NeuN but not OLIG2, SOX10, PDGFRA or NG2 in the cortex. In contrast, the tdT^+^ cells expressed all of these markers (OLIG2, SOX10, PDGFRA, NG2 and NeuN) in the striatum. (F) The ratio of the OLIG2^+^ and SOX10^+^ cells of the electroporated cells in the cortex. N = 4 mice per group. (G) Fate mapping of LGE-derived cells at P0. tdT^+^ cells expressed SOX10 and SOX9 in the striatum at P49.

To further determine whether LGE-derived OPCs migrate into the cortex after E18.5, we used the same strategy as mentioned above (Figure 1-figure supplement 1A). We observed that at P10, the majority of the tdT^+^ cells were located in the striatum, whereas very few tdT^+^ cells were located in the neocortex (Figure 1-figure supplement 1B). All tdT^+^ cells in the neocortex expressed NeuN but did not express OLIG2 (Figure 1-figure supplement 1C). In fact, tdT^+^ cells in the neocortex did not express OPC marker genes; instead, they expressed NeuN in the neocortex. In contrast, tdT^+^ cells in the striatum expressed OPC marker genes (*Olig2*, *Sox10* and *Pdgfra*) and NeuN (Figure 1-figure supplement 1D), consistent with our observation at E18.5 (Figure 1E). We also found that tdT^+^ cells expressed the astrocyte marker ALDH1L1 in the striatum at P10 (Figure 1-figure supplement 1D). Furthermore, through consecutive tissue sectioning and 3D reconstruction, we revealed that the traced cells tended to be localized in the striatum (Figure 1-figure supplement 1F). Collectively, our findings indicated that the LGE gives rise to striatal OLs, neurons and astrocytes, but not cortical OLs.

### Lineage tracing of LGE-derived OPCs by combining IUE with the PiggyBac transposon system

To further investigate whether the LGE generates cortical OPCs during cortical development, we employed an alternative fate mapping strategy by combining IUE with the PiggyBac transposon system to map the fate of OPCs derived from LGE. This approach utilized a transposase enzyme (helper plasmid) to integrate the H2B-GFP sequence from a donor plasmid into the genome of the in utero electroporated LGE progenitors along the lateral ventricular zone. The H2B-GFP reporter vector was delivered specifically into the LGE VZ of wild-type embryos at E13.5 by IUE, and these embryos were examined at E18.5 or P10 (Figures 2A-B). In this system, most traced cells (GFP^+^) were located in the striatum, and very few traced cells were observed in the cortex (Figures 2C and E). Usually, a small population of RGCs located in the dorsal LGE or lateral/ventral pallium was labeled by IUE. These RGCs mainly migrated to the lateral/ventral pallium and amygdala (Figure 2C). We analyzed 139 GFP^+^ cells in the cortex (N = 4), but we did not observe any GFP^+^ cells that were OPCs in the dorsal cortex, as these cells did not express OLIG2, SOX10 or PDGFRA (Figure 2D). Instead, they expressed NeuN (Figure 2D). In contrast, GFP^+^ cells expressed OLIG2, SOX10, PDGFRA and NeuN in the striatum at E18.5 (Figure 2D). The same results were also observed at P10 (Figure 2E). Again, our findings strongly suggested that the LGE may not generate cortical OPCs.

**Figure 2.**
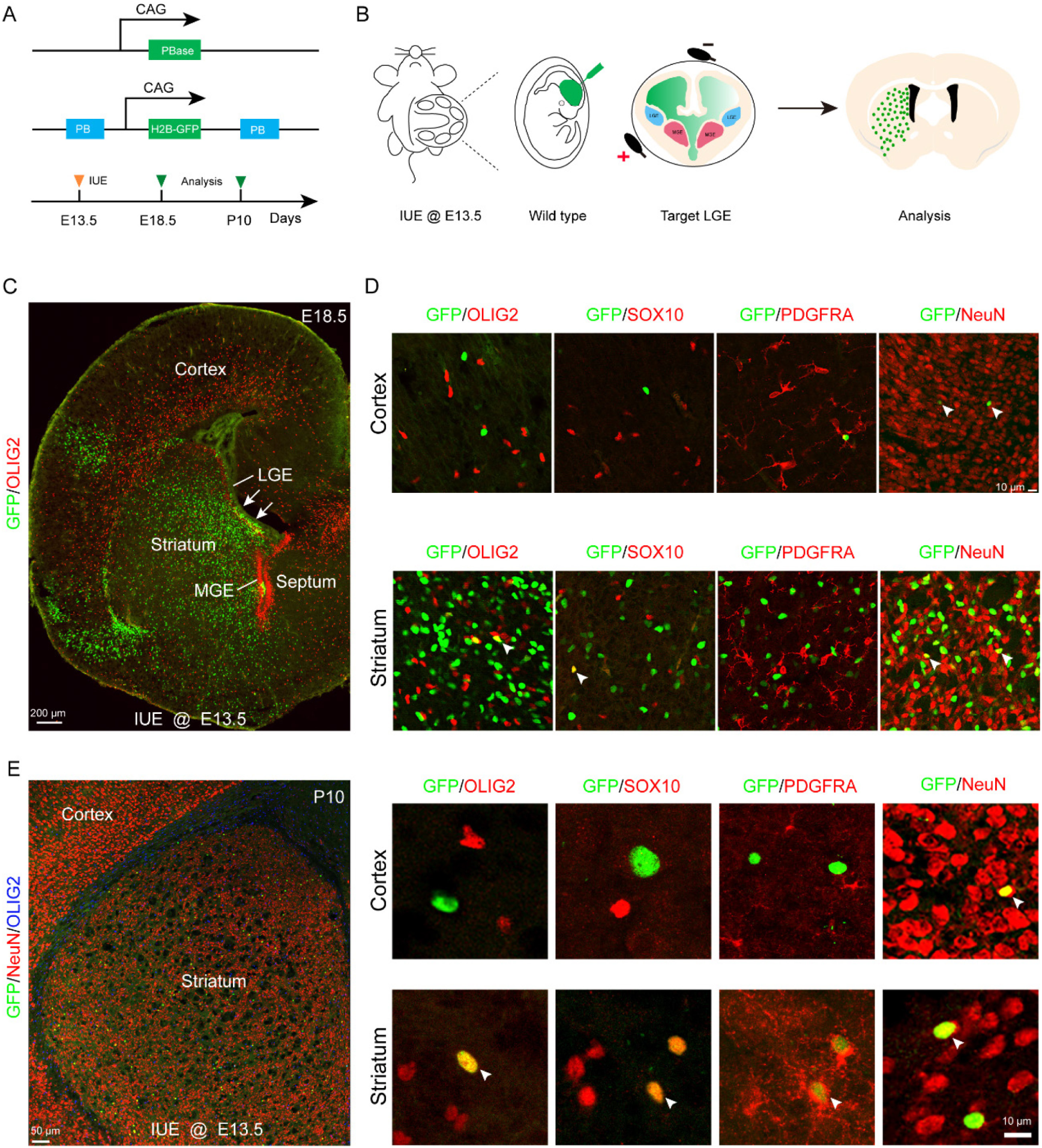
Fate mapping of LGE-derived OPCs by combining IUE with PiggyBac transposon system. (A) Schematic of the PiggyBac transposon reporter system used in this study. (B) The experimental workflow. (C) Representative coronal sections showing the distribution of GFP^+^ cells at E18.5. (D) GFP^+^ cells expressed NeuN but not glial cell markers, such as OLIG2, SOX10 and PDGFRA in the cortex. However, GFP^+^ cells expressed NeuN, OLIG2, SOX10 and PDGFRA in the striatum. (E) GFP^+^ cells expressed NeuN but not glial cell markers, such as OLIG2, SOX10 and PDGFRA in the cortex and GFP^+^ cells expressed NeuN, OLIG2, SOX10 and PDGFRA in the striatum at P10.

### An exclusion strategy revealed that LGE/CGE contribution to cortical OPCs is minimal

A previous report^9^ suggested that cortical OPCs originate from MGE, LGE/CGE and cortical RGCs. When MGE-derived cortical OPCs were eliminated, LGE/CGE-derived cells accounted for approximately 50% of cortical OPCs at P10. Subsequently, another study reported that approximately 20% OLs in corpus callosum are ventral-derived at P12/P13^26^. However, according to our recent findings^10,11^, we speculate that these OPCs (*Gsx2^+^* lineage) may derived from the cortical progenitors (*Emx1^+^* lineage). To confirm these findings, using the *Emx1^Cre^;* H2B-GFP mice, we found that many GSX2^+^ cells were located in the cortical SVZ. As expected, these GSX2^+^ cells were co-labelled with GFP (*Emx1^+^* lineage), suggesting that cortical SVZ GSX2^+^ cells were derived from cortical progenitors (Figure 3-figure supplement 1A-C).

While distinct fate mapping strategies provided evidence that LGE may not generate cortical OPCs during cortical development, there remained a possibility that a small population of LGE/CGE RGCs or progenitors might generate a subpopulation of cortical OPCs at specific time points. To address this possibility, we crossed the *Emx1*^*Cre* 27–29^, *Nkx2.1*^*Cre* 30^ and *Rosa^H2B-GFP^*^18^ mouse lines to generate *Emx1^Cre^; Nkx2.1^Cre^; Rosa^H2B-GFP^* mice. Because *Emx1^Cre^* was specifically expressed in cortical RGCs^27^ and *Nkx2.1^Cre^* was specifically expressed in MGE/AEP RGCs^30^, cortical OPCs were traced (GFP^+^ cells) in using *Emx1^Cre^; Nkx2.1^Cre^; Rosa^H2B-GFP^* mice. GFP^-^ cell OPCs, on the other hand, were considered as LGE/CGE-derived cortical OPCs (OLs) (Figures 3A-B). We performed double staining of GFP with OPC marker genes, such as *Sox10* and *Pdgfra* in *Emx1^Cre^; Nkx2.1^Cre^;* H2B-GFP mice (Figure 3C). We observed that the majority of OPCs or OLs were co-labeled with GFP in *Emx1^Cre^; Nkx2.1^Cre^; Rosa^H2B-GFP^* mice at P10 (Figure 3C). Statistical analysis showed that only approximately 2% of OPCs were GFP^-^ cells, and they were possibly derived from the LGE/CGE (Figures 3D-E), which was consistent with our observation at P0 (Figures 3G-I). Importantly, the expression of *Nkx2.1^Cre^* is lower within the dorsalmost region of the MGE than in other *Nkx2.1*-expressing regions^30^. Thus, these GFP^-^ OPCs (∼2%) may be derived from the dorsal MGE rather than from the LGE/CGE. Using this approach, we revealed that nearly all cortical OLs/OPCs were derived from MGE/AEP and cortical RGCs. Taken together, our findings indicated that the contribution of cortical OPCs from the LGE/CGE is minimal.

**Figure 3.**
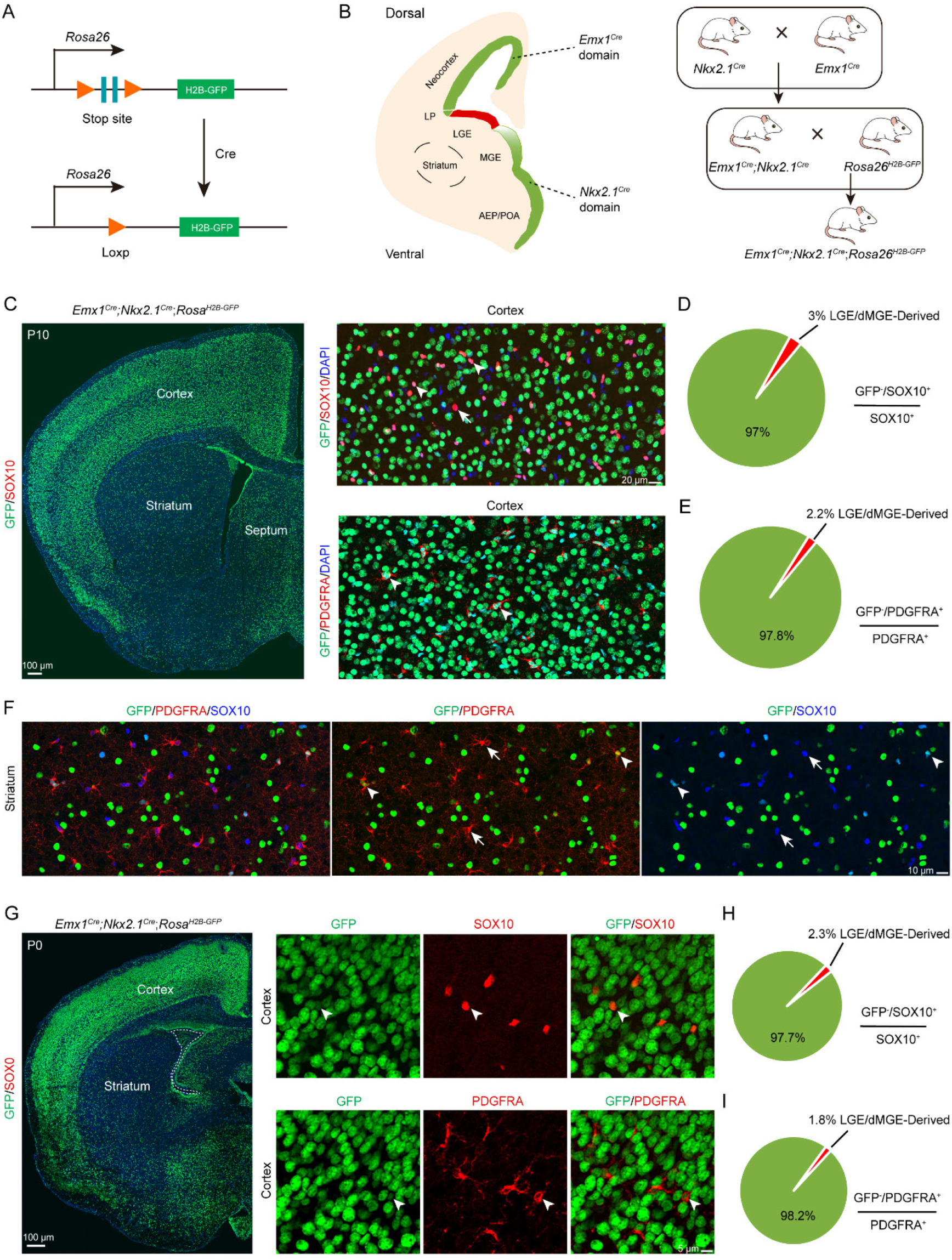
An exclusion strategy showed that LGE/CGE contribution to cortical OPCs is minimal. (A) Scheme of the H2B-GFP reporter lines. (B) Experimental design of the exclusion strategy to trace the lineage of LGE RGCs. (C) Representative coronal sections showing the traced cells in the forebrain. The majority of SOX10- and PDGFRA-positive cells were GFP^+^ cells in the cortex at P10. (D-E) The pie chart shows that the percentage of LGE/dMGE-derived cortical OPCs was approximately 3%. N = 6 mice per group. (F) Many GFP^+^ cells expressed SOX10 and PDGFRA in the striatum at P10. (G) Nearly all SOX10 and PDGFRA expressed GFP in the cortex at P0. (H-I) The pie chart shows that the percentage of LGE/dMGE-derived cortical OPCs was less than 3%. N = 6 mice per group.

### The MGE/AEP was the predominant ventral source of the cortical OPCs

*Olig2*, a basic helix-loop-helix transcription factor, plays a critical role in the regulation of OPC fate determination. To investigate the role of *Olig2* in the MGE, we crossed *Nkx2.1^Cre^* mice with *Olig2^F/+^* mice to eliminate *Olig2* expression in the SVZ/VZ of the MGE/AEP (Figures 4A-B). We referred to the conditional mutants as *Olig2-*NCKO mice. A previous study^9^ showed that MGE/AEP-derived OPCs migrate into the cortex around E14.5, and LGE/CGE-derived OPCs migrate into cortex around E15.5. We assumed that the MGE/AEP was the predominant ventral source of cortical OPCs, and that inhibiting MGE/AEP-derived OPC generation would therefore significantly reduce the number of cortical OPCs at E16.5. To test this possibility, we analyzed *Olig2*-NCKO mice, in which the generation of MGE/AEP-derived OPCs was inhibited. Our results showed that the expression of OPC marker genes, such as *Sox10* and *Pdgfra* was significantly reduced (> 10-fold) in the cortex of *Olig2*-NCKO mice, compared to the control mice at E14.5 and E16.5 (Figures 4C-E). Considering the recombination efficiency of the CRE enzyme, the limited presence of residual OPCs in the cortex of *Olig2*-NCKO mice may be arise from the MGE/AEP. Indeed, we observed few PDGFRA-positive cells in the MGE/AEP of the *Olig2*-NCKO mice (Figure 4F). Altogether, our findings indicated that the MGE/AEP was the predominant ventral source of cortical OPCs.

**Figure 4.**
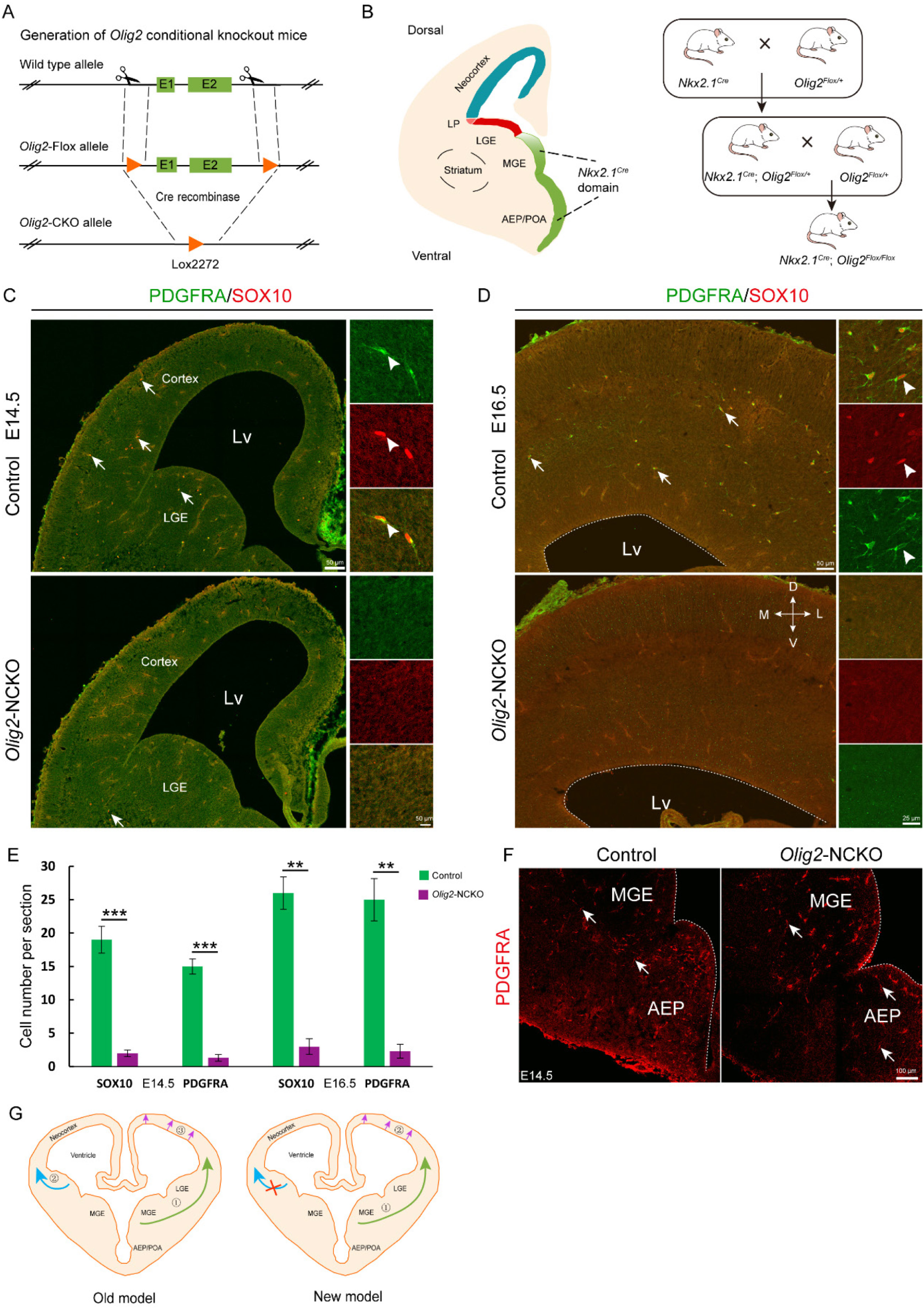
The MGE may be the sole ventral source of cortical OPCs. (A) The CRISPR/Cas9 technique was used to generate an *Olig2* conditional knockout allele. (B) Experimental design for the generation of *Olig2*-NCKO mice. (C-D) The expression of SOX10 and PDGFRA was significantly reduced in the cortex of *Olig2*-NCKO mice compared with that of control mice. (E) The number of SOX10- and PDGFRA-positive cells was significantly reduced in *Olig2*-NCKO mice compared with control mice. Student’s t-test, **P <.01, ***P <.001, n ≥ 5 mice per group, mean ± SEM. (F) Few PDGFRA-positive cells were detected in the MGE/AEP of the *Olig2*-NCKO mice. (G) A new model of the developmental origins of cortical OPCs.

## Discussion

It is widely accepted in the field of developmental neuroscience that cortical OPCs originate from three regions: the MGE/AEP, LGE/CGE and cortex. However, in the current study, we re-examined the developmental origins of cortical OPCs using diverse strategies. Our findings show that cortical OPCs may not come from the LGE/CGE, contrary to previous reports^9,12^, which suggested that the LGE/CGE generates cortical OPCs (Figure 4G). In fact, no cortical OPCs migrating from the LGE to the cortex were observed via our two fate mapping strategies. Furthermore, an exclusion strategy suggested that limited contribution of the LGE/CGE to cortical OPCs. In addition, according to our study of *Olig2*-NCKO mice, MGE/AEP seems to be the predominant ventral source of cortical OPCs during neocortical development. Overall, these findings provide strongly evidence that the contribution of the cortical OPCs from the LGE/CGE is minimal in the mouse brain.

### Why LGE is not the main OPC pool for the cortex during development

OLs are specialized cells of the CNS that produce myelin, a multilayered membrane that spirally enwraps axons and facilitates rapid nerve conduction. Recent studies have shown that cortical OLs are heterogeneous, as there are transcriptionally distinct subpopulations of cortical OLs during development and disease^8,31–33^. Indeed, OLs are derived from distinct OPC pools, which acquire different abilities to respond to demyelination^9,34^. In the spinal cord, the developmental origin of OLs has been hotly debated for many years. Some groups believed that OLs originate only from the ventral neural tube, whereas others tend to support multiple origins^12,35–37^. One pioneering study^9^ purported that, in the forebrain, cortical OLs derive from multiple origins, such as the MGE, LGE and cortex.

In general, the specification of OLs depends on SHH protein released from the notochord and floor plate, as *Shh* expression is both necessary and sufficient for OL generation in the spinal cord^38^. As in the spinal cord, OL generation in the cortex is also dependent on the *Shh* signaling pathway^39^. During MGE development, *Nkx2.1* and *Six3* promote SHH secretion. Furthermore, the early-born neurons of the MGE also secrete SHH protein before E14.5^40^. Thus, the high levels of SHH in the MGE contribute to the generation of OPCs from RGCs. In contrast, *Shh* signaling pathway activity is very weak in the LGE at the early stage (before E14.5). As the level of SHH gradually increases in the LGE, this results in a shift from neurogenesis to gliogenesis after E15.5, and the large number of the striatal neurons may block the migration of OPCs into the neocortex^41,42^. In summary, the process of cortical OL generation can be summarized as follows: First, the high level of SHH in the MGE allows for the production of a small population of cortical OPCs around E12.5. Subsequently, multipotent intermediate progenitors begin to express DLX transcription factors resulting in ending the generation of OPCs in the MGE^43–45^. Second, radial glial cells (RGCs) undergo a gradual transition from neurogenesis to gliogenesis around E15.5, driven by the progressive elevation of SHH levels in the LGE. However, the LGE had already generated a substantial number of neurons during this period, blocking the migration of OPCs into the cortex. Finally, with the gradual increase of SHH concentration, cortical RGCs begin to cease neurogenesis and shift toward gliogenesis after E16.5^10,41,46^. Therefore, the generation of cortical OPCs is regulated by both precise temporal and spatial patterning, which is strictly dependent on the local environment.

### Cortical-derived OLs/OPCs make up the majority of the cortical OLs/OPCs in the adult brain

During the early development of the cortex, both pallium and subpallium can generate cortical OPCs^9,47^. Interestingly, most of the ventral-derived OPCs, particularly MGE-derived OPCs/OLs, are eventually eliminated due to the continued expansion of dorsal-derived OPCs in the adult cortex^6,9^. It appears that OPCs generated later in development may possess a competitive advantage over their earlier-generated counterparts. Indeed, human glial progenitor cells (hGPCs) engrafted neonatally or transplanted into the adult mouse brain can effectively disperse throughout the forebrain, differentiate into mature OLs and remyelinate the demyelinated adult brain^1,48^. These findings indicate that OPCs generated later in development have the ability to take over for earlier-generated OPCs in the adult brain.

Based on the present work and other studies^8,9,47^, cortical-derived OPCs probably constitute the majority of cortical OPCs, including mature cortical OLs. Thus, OLs may function as a homogeneous population in the adult forebrain. In fact, the number of LGE/CGE-derived cortical OPCs may also gradually decrease in the adult brain, as reported by Lui et al^6^. However, Lui et al. did not analyze LGE-derived and MGE-derived OPCs separately. Therefore, they did not exclude the possibility that the LGE/CGE does not generate cortical OPCs. In summary, cortical OPCs/OLs comprise MGE-derived and cortical RGC-derived OPCs/OLs during the neonatal stage. In contrast, cortical-derived OPCs/OLs make up the majority of the cortical OPCs/OLs in the adult cortex. Recently, another study showed that ventral-derived OLs contribution to adult cortex is minimal^49^.

In summary, our study provides strong evidence that the contribution of the LGE to cortical OPCs is minimal during forebrain development. Thus, our findings provide new evidence about the developmental origins of cortical OPCs, which may guide future research on both normal development and disease.

## Methods

### Animals

All experiments conducted in this study were in accordance with the guidelines of Fudan University. We used *Emx1*^*Cre* 27,28^, and *Nkx2.1^Cre^* ^30^ mice, in conjunction with either *Rosa^H2B-GFP^*^18^ or *Rosa-IS*^18^ for genetic fate mapping. *Olig2^F/+^* mice were generated via the CRISPR/Cas9 strategy. Loxp sites flanked the coding region of exons 1 and 2. Wild-type mice or littermates without the Cre allele were used as controls. All mice were maintained on a mixed genetic background (C57BL/6J and CD1). The mice were allowed access to water and food ad libitum and maintained on a 12-hour light/dark cycle. The day of vaginal plug detection was considered embryonic day 0.5 (E0.5), and the day of birth was defined as postnatal day 0 (P0). Both male and female mice were used in all experiments.

### Tissue preparation

Postnatal mice were deeply anesthetized and perfused intracardially thoroughly with ice-cold PBS followed by 4% paraformaldehyde (PFA). Embryos were harvested from deeply anesthetized pregnant mice. All brains were fixed overnight in 4% PFA at 4°C, cryoprotected in 30% sucrose for at least 24 hours, embedded in optimal cutting temperature (O.C.T.) (Sakura Finetek) on dry ice and ethanol slush and preserved at −80°C. All frozen mouse tissues were sectioned into 20 µm thick sections unless specifically mentioned and stained on glass slides.

### Plasmid Construction

The pCAG-Cre vector was constructed as reported elsewhere. The coding sequence of Cre in the pCAG-Cre vector was replaced by the PiggyBac transposase to construct the pCAG-PiggyBac transposase vector. The coding DNA sequence (CDS) of Cre in the pCAG-Cre vector was replaced by H2B-GFP from the *Rosa-*H2B-GFP mouse strain to construct the PB-pCAG-H2B-GFP vector. The inverted terminal repeat (ITR) sequence of the PiggyBac transposon was cloned upstream of the *CAG* promoter and downstream of the beta-globin polyA sequence for recognition by the PiggyBac transposase.

### In utero electroporation (IUE)

In utero electroporation (IUE) of wild-type or IS embryos was performed at E13.5. The pCAG-Cre (Addgene #13775), pCAG-GFP (Addgene #11150), pCAG-PiggyBac transposase or PB-pCAG-H2B-GFP (final concentration: 1-2 μg/μL) plasmid was mixed with 0.05% Fast Green (Sigma) and injected into the lateral ventricle of embryos (0.5 μL per embryo) using a beveled glass micropipette. Five electrical pulses (duration: 50 ms) were applied at 42 V across the uterine wall with a 950 ms interval between pulses. Electroporation was performed using a pair of 5 mm platinum electrodes (BTX, Tweezertrode 45-0488, Harvard Apparatus) connected to an electroporator (BTX, ECM830). The embryos were analyzed at E18.5 and P10.

### Immunofluorescence

The sections were first washed with 0.05 M TBS for 10 min, incubated in Triton-X-100 (0.5% in 0.05 M TBS) for 30 min at room temperature (RT), and then incubated with blocking solution (5% donkey serum +0.5% Triton-X-100 in 0.05 M TBS, pH= 7.2) for 2 h at RT. The primary antibodies were diluted in 5% donkey serum, incubated overnight at 4°C, and then rinsed 3 times with 0.05 M TBS.

Secondary antibodies (1:500, all from Jackson ImmunoResearch) matching the appropriate species were added and incubated for 2-4 h at RT. Fluorescently stained sections were then washed 3 times with 0.05 M TBS for 10 min. The sections were then stained with 4’,6-diamidino-2-phenylindole (DAPI) (1:3000, Sigma) diluted in TBS for 1 min and then finally rinsed three more times with TBS. The sections were then coverslipped with Gel/Mount (Biomeda, Foster City, CA). The primary antibodies used in this study are shown in the key resource table.

### *In situ* hybridization (ISH)

ISH was performed on 20-μm cryosections using digoxigenin riboprobes as previously described ^50^. The sections were first postfixed in 4% PFA for 20 min. mRNA *in situ* hybridization was performed as previously described^49^. The probes were constructed from cDNA isolated from the wild-type mouse brain and amplified by PCR using the following primers:

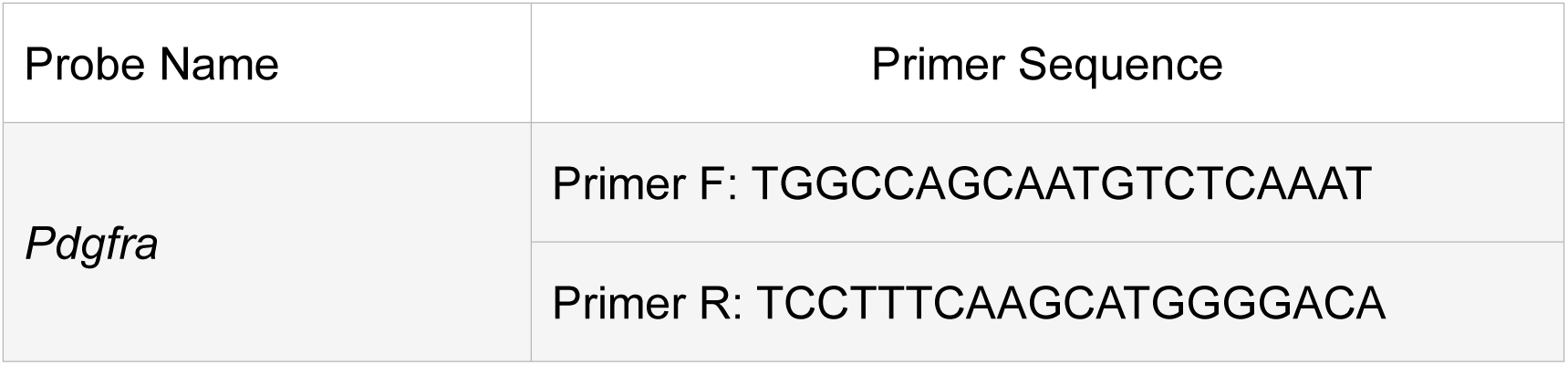

### 3D Reconstruction

Images of serial coronal sections along the rostral–caudal axis were collected via an Olympus VS 120 microscope using a 10X objective and used by Amira as a formwork for reconstruction. The coordinates of each labeled cell (tdT^+^) in the brain were determined. A stereotaxic mouse brain atlas was used to locate the brain anatomical structures where the labeled cells were positioned.

### Image acquisition and statistical analysis

Images for quantitative analyses were acquired with an Olympus VS 120 microscope using a 10X objective. Bright-field images were acquired with an Olympus VS 120 microscope using a 10X objective. Other fluorescence images were taken with the Olympus FV3000 confocal microscope system using 20× or 40× objectives. The images were merged, cropped and optimized in Adobe Photoshop and Adobe Illustrator without distorting the original visual information.

Analyses were performed using Microsoft Excel. Unpaired t test was used to determine statistical significance. All quantitative data are presented as the mean ± SEM. Differences with *p* values < 0.05 were considered significant. For quantification of SOX10^+^, PDGFRA^+^ and GFP^+^ cells in the cortex at E18.5 and P10, six anatomically matched 20-μm thick coronal sections were selected (n = 6 mice per group). We counted SOX10^+^, PDGFRA^+^ and GFP^+^ cells in the cortex under a 20X objective lens. The data are presented as the number of SOX10^+^, PDGFRA^+^ and GFP^+^ cells per section for each cortex.

For quantification of SOX10^+^ and PDGFRA^+^ cells in the dorsal cortex at E14.5 and E16.5, five or six anatomically matched 20-μm thick coronal sections were selected (n = 6, E14.5; n=5, E16.5). We counted SOX10^+^ and PDGFRA^+^ cells in the cortex under a 20X objective lens. The data are presented as the number of SOX10^+^ and PDGFRA^+^ cells per section for each cortex.

### Key resource table

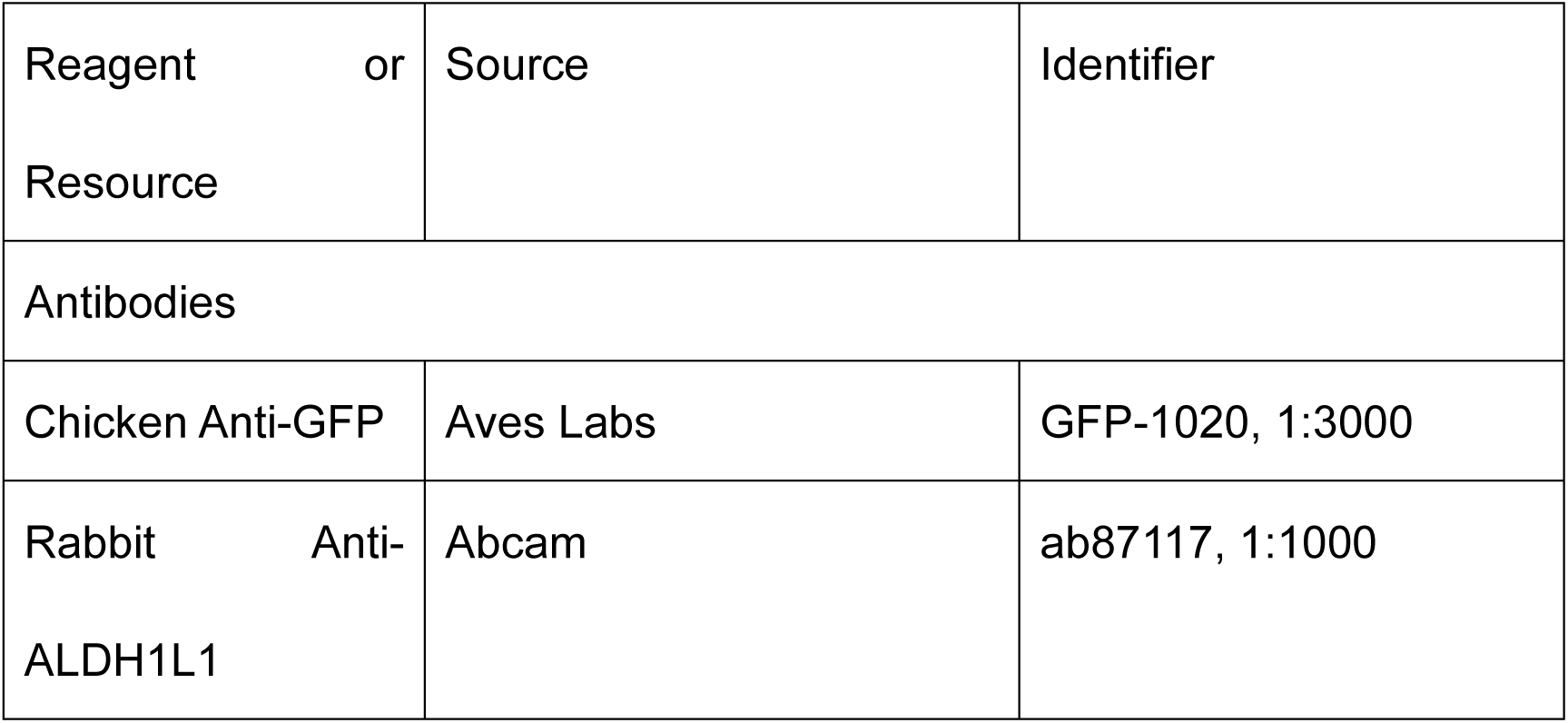
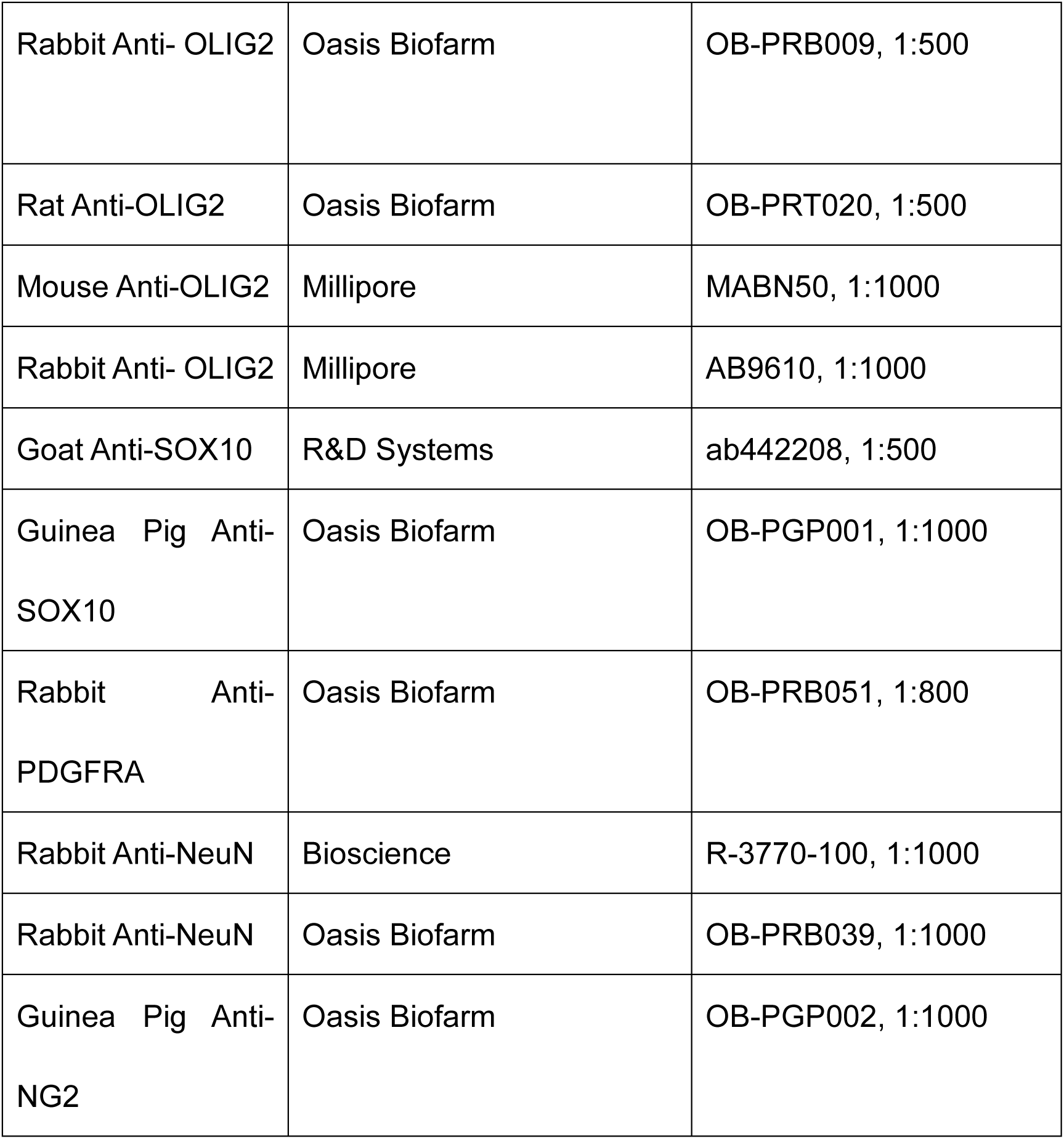

Experimental models

|  |  |  |
| --- | --- | --- |
| Mouse: C57BL/6 | Department of Laboratory<br>Animal Science at Fudan<br>University | http://10.107.12.196/ |
| Mouse: CD1 | Department of Laboratory<br>Animal Science at Fudan<br>University | http://10.107.12.196/ |
| Mouse: *IS* reporter | The Jackson Laboratory | Stock No. 028582 |
| Mouse: *Olig2<sup>F/+</sup>* | This manuscript | N/A; available from the authors |
| Mouse: *Nkx2.1<sup>Cre</sup>* | The Jackson Laboratory | Stock No. 008661 |
| Mouse: *Emx1<sup>Cre</sup>* | The Jackson Laboratory | Stock No. 005628 |

Recombinant DNA

|  |  |  |
| --- | --- | --- |
| pCAGIG | Addgene | Cat#11159 |
| pGL4.10 | Promega | Cat#E6651 |
| pGL4.23 | Promega | Cat#E8411 |

All secondary antibodies were from Jackson ImmunoResearch Labs.

| Secondary Antibodies | Cat Number | Dilution |
| --- | --- | --- |
| Alexa®488-Conjugated Affinipure Donkey Anti-Chicken IgY<sup>++</sup> (IgG) (H<sup>+</sup>L) | 703-545-155 | 1:500 |
| Cy<sup>TM</sup>3-Conjugated Affinipure Donkey Anti-Rabbit IgG (H<sup>+</sup>L) | 711-165-152 | 1:500 |
| Alexa®647-Conjugated Affinipure Donkey Anti-Rat IgG (H+L) | 712-605-153 | 1:500 |
| Alexa®647-Conjugated Affinipure Donkey Anti-Goat IgG (H<sup>+</sup>L) | 705-605-147 | 1:500 |
| Alexa®488-Conjugated Affinipure Donkey Anti-Rabbit IgG (H<sup>+</sup>L) | 711-545-152 | 1:500 |
| Cy™3-AffiniPure Donkey Anti-Goat IgG (H<sup>+</sup>L) | 705-505-147 | 1:500 |
| Cy™3-AffiniPure Donkey Anti-Rat IgG (H<sup>+</sup>L) | 712-165-150 | 1:500 |
| Cy™3-AffiniPure Donkey Anti-Guinea Pig IgG (H<sup>+</sup>L) | 706-165-148 | 1:500 |

## Acknowledgements

This work was supported by the Ministry of Science and Technology of China (STI2030-2021ZD0202300), National Natural Science Foundation of China (NSFC 31820103006, 82271197, 81974175 and 32200776), Shanghai Municipal Science and Technology Major Project (No. 2018SHZDZX01), ZJ Lab, and Shanghai Center for Brain Science and Brain-Inspired Technology.

## Data availability statement

In this study, no dataset was utilized. However, any supplementary information necessary for the reanalysis of the data presented in this paper is obtainable from the corresponding author upon request.

