## Supplemental Figures for "The Lateral/Caudal Ganglionic Eminence Makes a Limited Contribution to Cortical Oligodendrocytes": Supplement.pdf

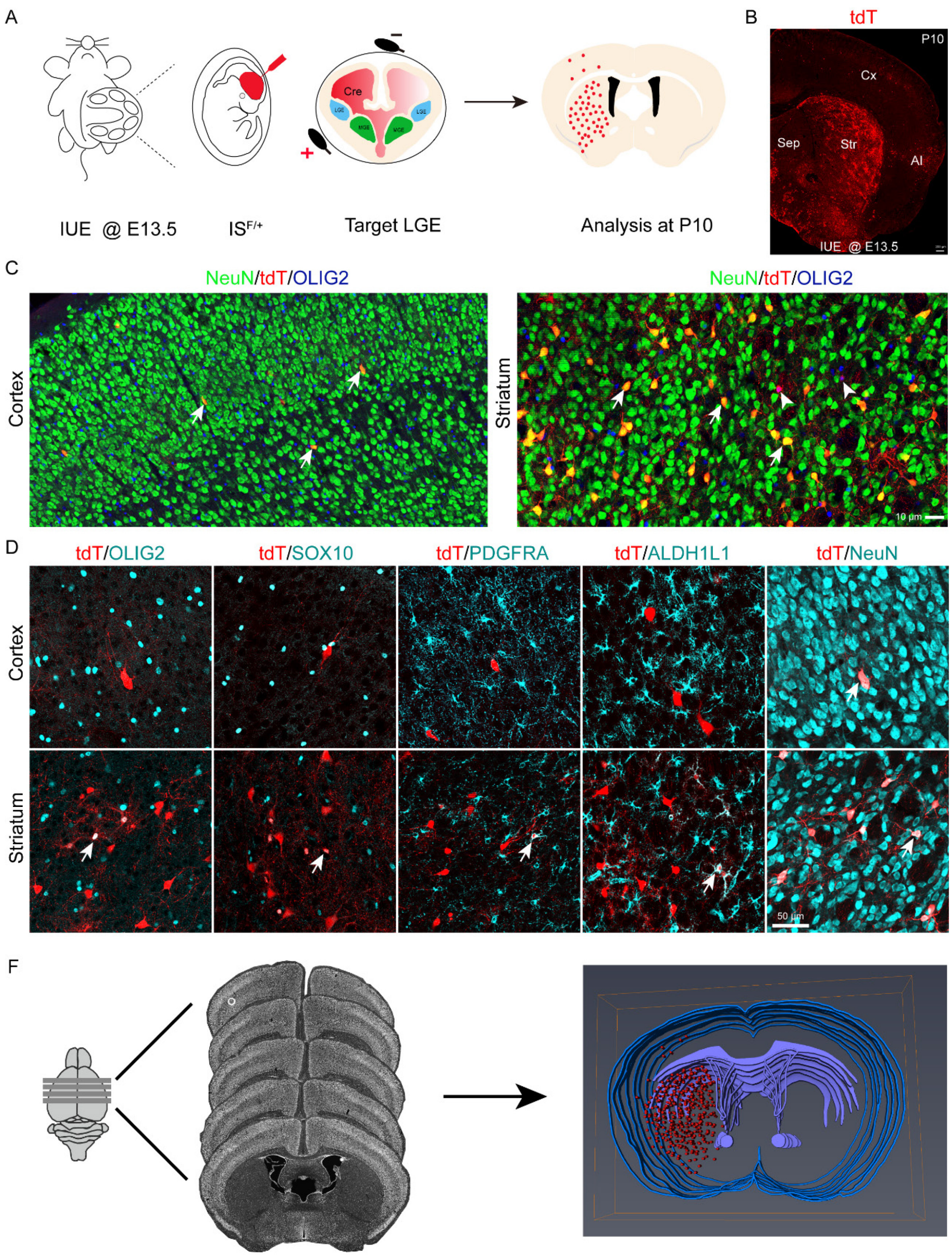


**Figure1-supplement 1. Lineage tracing of LGE-derived OPCs by combining IUE with a Cre recombinase-dependent IS reporter**

(A) Experimental schedule for fate mapping of LGE-derived OPCs.

(B) Representative coronal sections showing the distribution of the tdT$^{+}$ cells at

P10.

(C) Nearly all $tdT^+$ cells expressed NeuN but not OLIG2 in the cortex. $tdT^+$ cells expressed NeuN and OLIG2 in the striatum.

(D) $tdT^+$ cells did not express OLIG2, SOX10, PDGFRA or ALDH1L1. Instead, they expressed NeuN in the cortex. In sharp contrast, $tdT^+$ cells in the striatum expressed OLIG2, SOX10, PDGFRA, ALDH1L1 and NeuN at P10.

(E) 3D reconstruction of consecutive brain sections demonstrated that the traced cells were mainly located in the striatum.

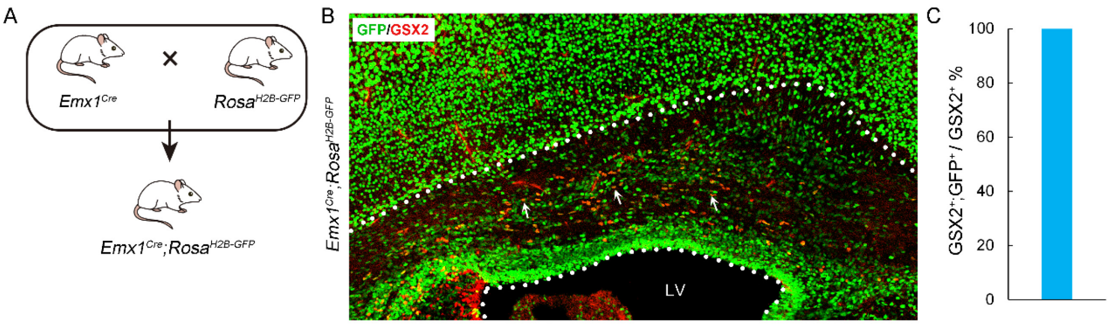


**Figure3-supplement 1. Lineage tracing of *Emx1*$^{Cre}$ derived cortical cells.**

(A) Experimental design for the generation of *Emx1*$^{Cre}$; H2B-GFP mice.

(B) Double immunostaining of GSX2 with GFP in the *Emx1*$^{Cre}$; H2B-GFP cortex at P0.

(B) The statistics show that nearly all $GSX2^+$ cells co-labelled with GFP in the cortical SVZ.
